# Sensitivity to Envelope Interaural Time Difference: Models of Diverse LSO Neurons

**DOI:** 10.1101/2020.09.08.288282

**Authors:** Andrew Brughera, Jimena A. Ballestero, David McAlpine

## Abstract

A potential auditory spatial cue, the envelope interaural time difference (ITD_ENV_) is encoded in the lateral superior olive (LSO) of the brainstem. Here, we explore computationally modeled LSO neurons, in reflecting behavioral sensitivity to ITD_ENV_. Transposed tones (half-wave rectified low-frequency tones, frequency-limited, then multiplying a high-frequency carrier) stimulate a bilateral auditory-periphery model driving each model LSO neuron, where electrical membrane impedance low-pass filters the inputs driven by amplitude-modulated sound, limiting the upper modulation rate for ITD_ENV_ sensitivity. Just-noticeable differences in ITD_ENV_ for model LSO neuronal populations, each distinct to reflect the LSO range in membrane frequency response, collectively reproduce the largest variation in ITD_ENV_ sensitivity across human listeners. At each stimulus carrier frequency (4-10 kHz) and modulation rate (32-800 Hz), the top-performing model population generally reflects top-range human performance. Model neurons of each speed are the top performers for a particular range of modulation rate. Off-frequency listening extends model ITD_ENV_ sensitivity above 500-Hz modulation, as sensitivity decreases with increasing modulation rate. With increasing carrier frequency, the combination of decreased top membrane speed and decreased number of model neurons capture decreasing human sensitivity to ITD_ENV_.

## Introduction

Human listeners exploit differences in the intensity and timing of sounds arriving at the two ears—interaural intensity differences (IIDs) and interaural time differences (ITDs), respectively—to determine the location of the source on the horizontal plane (Blauert, 1997). For sound frequencies below about 1400 Hz, human IIDs are small except for sources very close to the head (Brungart & Rabinowitz, 1999), and source location is determined primarily from ITDs conveyed in the temporal fine structure (TFS, ITD_TFS_) of sounds (Wightman & Kistler, 1992). At higher sound frequencies, listeners exploit IIDs for source localization (Mills, 1960; Sandel et al., 1955) and, under laboratory conditions at least, are sensitive to ITDs conveyed in the modulated envelope of sound pressure, referred to as “envelope ITD” (ITD_ENV_) (McFadden & Pasanen, 1976). Human abilities to exploit ITD_ENV_ under natural listening conditions remains an open question, which is particularly relevant to listeners with bilateral cochlear implants (bCIs, CIs): as clinical CI processors convey only the correct envelope, and not the correct TFS (Gransier et al., 2020).

Sensitivity to binaural spatial cues relies on neural encoding in the medial and lateral superior olive (MSO and LSO, respectively) of the auditory brainstem. MSO neurons, innervated with bilateral excitatory and inhibitory inputs, encode ITD_TFS_ in low-frequency sounds (Goldberg & Brown, 1968, 1969; Grothe & Sanes, 1993; Yin & Chan, 1990), by mechanisms of fast binaural-coincidence detection temporally sharpened by low-threshold potassium (K_LT_) channels (Mathews et al., 2010). LSO neurons, innervated by ipsilateral excitatory and contralateral inhibitory inputs (Fig. 1A), encode IID (Boudreau & Tsuchitani, 1967) and ITD_ENV_ conveyed in high-frequency sounds (Batra et al., 1997; Joris, 1996; Joris & Yin, 1995, 1998). Some LSO neurons express K_LT_ channels in rapidly-responding membranes, while others, increasingly common in higher-frequency regions of the LSO, have more slowly-responding integrative membranes, consistent with the encoding of IID (Barnes-Davies et al., 2004; Remme et al., 2014).

**Fig. 1.**
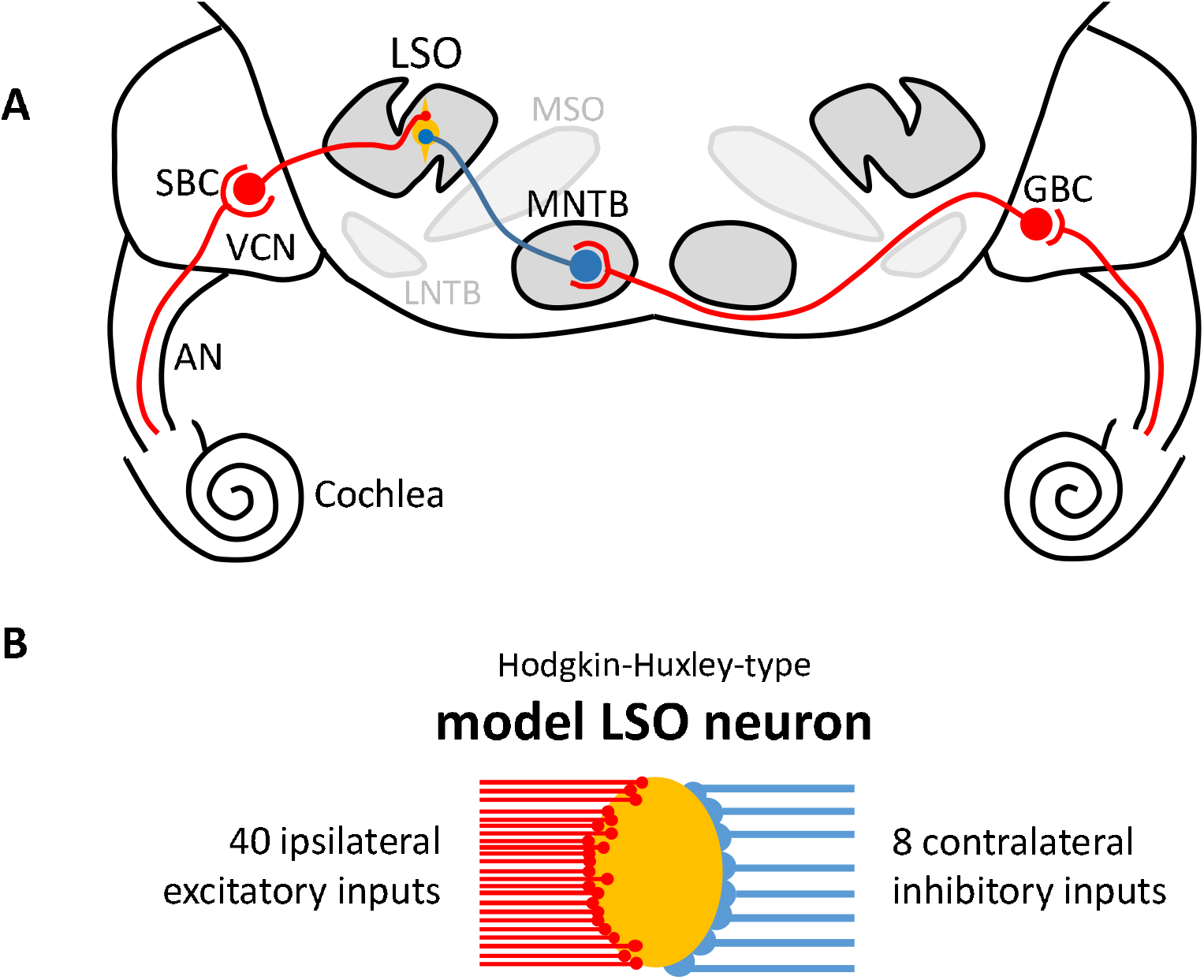
**A:** Inputs to the lateral superior olive (LSO). Excitatory neurons, axons, and synapses are in red; inhibitory in blue. The LSO receives ipsilateral excitation from spherical bushy cells (SBCs) of the ventral cochlear nucleus (VCN), each driven by 1-3 auditory-nerve (AN) fibers (ANFs) (Doucet & Ryugo, 2003; Lorente De No, 1981; Smith et al., 1993). The LSO receives contralateral inhibition from principal neurons of the medial nucleus of the trapezoid body (MNTB), each driven by a globular bushy cell (GBC) of the VCN; GBCs are driven by 9-70 ANFs (Banks & Smith, 1992; Glendenning et al., 1985; Smith et al., 1991; Spirou et al., 2005). **B:** Model LSO neuron with a single HH-type compartment, receiving simplified inputs directly from model ANFs: 40 ipsilateral excitatory inputs and 8 contralateral inhibitory inputs

Psychoacoustic studies show smaller just-noticeable differences (JNDs) in the ITD_TFS_ of low-frequency tones (Brughera et al., 2013), compared with larger JNDs in the ITD_ENV_ of modulated high-frequency tones (Bernstein & Trahiotis, 2002, 2014; Monaghan et al., 2015). These studies demonstrate two additional features specific to ITD_ENV_. First, sensitivity to ITD_ENV_ is extremely variable across listeners, including listeners highly trained in ITD_ENV_ discrimination, and exquisitely sensitive to ITD_TFS_: at each carrier frequency, ITD_ENV_ discrimination thresholds show four-to-ten-fold differences for the same modulation rate, and two-to-five-fold differences in the highest modulation rate with a measurable threshold. Second, with increasing carrier frequency, there is a decrease in ITD_ENV_ sensitivity, and in the maximum modulation rate with a measurable threshold. This worsening performance is paradoxical, as cochlear filters broaden in absolute bandwidth with increasing sound frequency (Glasberg & Moore, 1990), thereby better passing the modulation sidebands of amplitude-modulated sounds. This paradoxical decrease in ITD_ENV_ sensitivity with increasing carrier frequency, and the variance in ITD_ENV_ sensitivity across human listeners, remain unexplained.

Here, we explore computationally modeled LSO neurons, in reflecting behavioral sensitivity to ITD_ENV_. We extend a Hodgkin-Huxley-type model LSO neuron (Wang & Colburn, 2012) to populations, each distinct to reflect the range of membrane frequency responses observed in the LSO (Remme et al., 2014), with a membrane time constant from slow to very fast. Bilaterally innervating each model LSO neuron (Fig. 1B), an auditory-periphery model (Zilany et al., 2014) is stimulated by transposed tones (Fig. 2): high-frequency tones modulated with an envelope designed to evoke auditory temporal resolution reflecting TFS in low-frequency sounds (van de Par & Kohlrausch, 1997). At each stimulus carrier frequency and modulation rate, the top-performing model population generally reflects the best human performance. Model neurons of each speed provide the best performance over a particular range of modulation rate. ‘Off-frequency listening’ extends ITD_ENV_ sensitivity up to the modulation rates observed in the best-performing human listeners. Decreasing ITD_ENV_ sensitivity with increasing carrier frequency required reduced top model membrane speed and number of model neurons. The largest variation in sensitivity to ITD_ENV_ across human listeners is captured by the heterogeneity in model membranes.

**Fig. 2.**
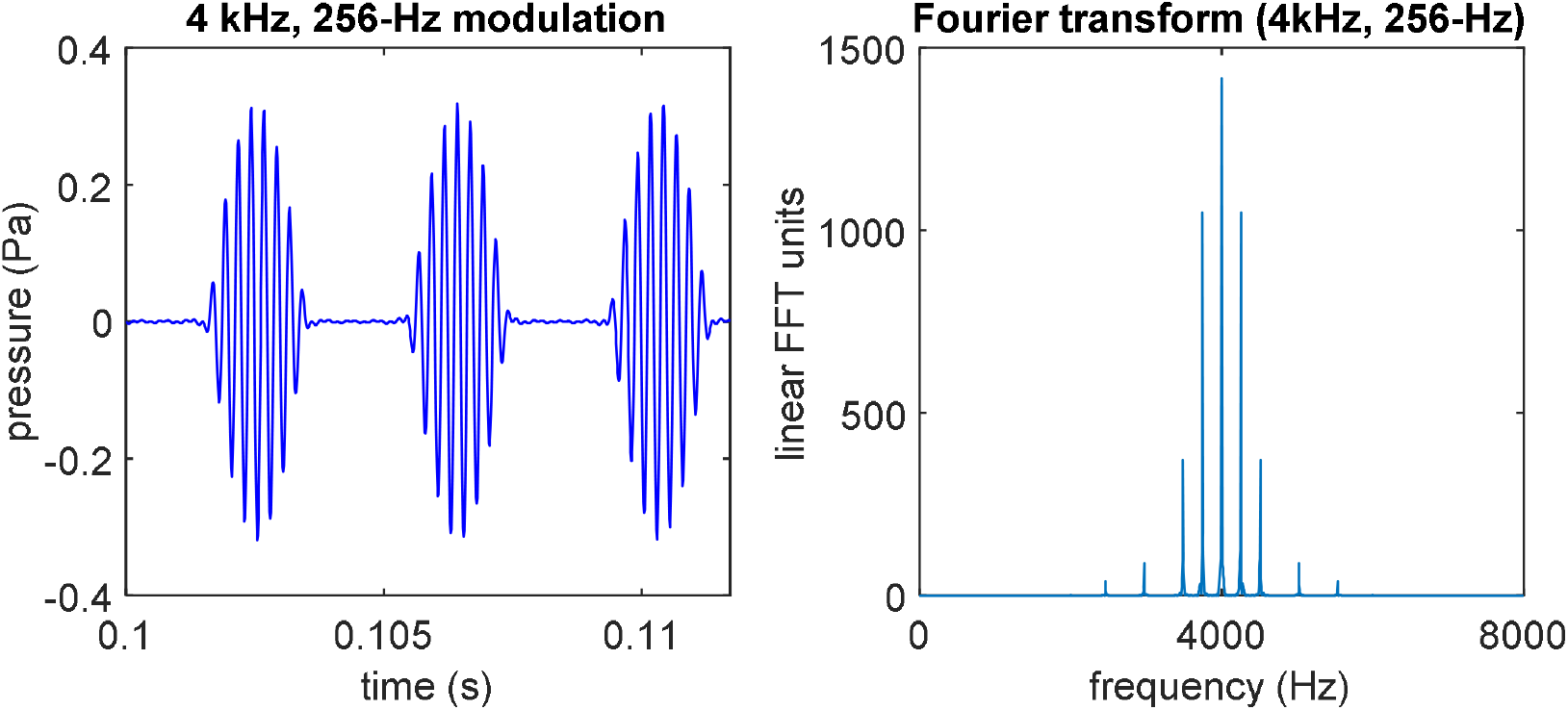
Transposed tone: a half-wave rectified low-frequency sinusoid, frequency-limited, then multiplying a high-frequency sinusoid as the carrier. Left: A transposed tone with a 4-kHz carrier, and modulation rate of 256 Hz. The envelope signal (not shown) is a 256-Hz sinusoid, half-wave rectified, with frequency components above 2 kHz then removed (leaving 0 Hz, 256 Hz, 512, 1024, and 1536 Hz, which includes even harmonics of 256 Hz). Forming the transposed tone, this envelope signal multiplies a 4-kHz sinusoidal carrier. Right: The fast Fourier transform of the transposed tone shows its frequency spectrum: a center peak at the 4-kHz carrier frequency, and modulation sidebands from the envelope signal, above and below the carrier by 256 Hz and its even harmonics up to 1536 Hz, with the extreme sidebands at 2464 and 5536 Hz. The limited frequency range of the envelope insures that this transposed tone has no frequency component below 2 kHz

## Methods

### Auditory Stimuli

The auditory stimuli were transposed tones (Fig. 2) (Bernstein & Trahiotis, 2002; van de Par & Kohlrausch, 1997): low-frequency tones, half-wave rectified, then band limited to frequencies below 2000 Hz, each multiplying a high-frequency carrier. Transposed-tone stimuli had maximum sound pressure level (SPL) 75 dB RMS, 300-ms duration, 20-ms cosine-squared ramps at onset and offset, carrier frequencies of 4, 6, and 10 kHz, and modulation rates of 32, 64, 128, 256, 512, and 800 Hz. Within each stimulus, phase of the carrier and envelope were steady. Across stimulus presentations, carrier phase was randomized, and the envelope phase (phase relative to the amplitude-modulation cycle) ranged from 0 to −525° in increments of 15° (negative phase is delay).

### Auditory-Periphery Model

Twenty-five repetitions of each transposed tone were applied bilaterally to an auditory-periphery model for humans (Glasberg & Moore, 1990; Zilany et al., 2014). With these stimulus repetitions, the stochastic auditory-periphery model provided 25 independent sets of spike times at each of the 36 envelope phases, for each of 40 model auditory-nerve (AN) fibers (ANFs) acting as excitatory synaptic inputs, and 8 model ANFs acting as inhibitory synaptic inputs, to the model LSO neurons (Fig. 1). Numbers of inputs are based on a recent model from anatomical counts (Gjoni et al., 2018). On-frequency listening conditions were implemented with ANFs having characteristic frequency (CF, the frequency of lowest spike-threshold in SPL) equal to the stimulus carrier frequency. Off-frequency listening conditions were implemented with ANFs having a single CF between the carrier frequency of the stimulus, and its first upper modulation sideband. Off-frequency listening in moderate, fast and very fast model LSO neurons separately applied ANFs with CF equal to 4.28, 4.43. 6.28, 6.43, 10.28, 10.43, or 10.56 kHz; HL-quick model neurons (named for their high conductances for h channels and leakage channels, and quick model membrane reflecting fast non-resonant LSO neurons) applied ANFs with CF equal to 4.28, 6.28, or 10.28 kHz. Model results for off-frequency listening are shown for CF equal to only 4.28, 6.28, and 10.28 kHz, providing a fair comparison with on-frequency listening. Simplified inputs to the model LSO (Fig. 1B) maintain the ipsilateral excitation and contralateral inhibition of the LSO, with the cochlear nuclei and trapezoid body modeled as relays without delay.

### Model LSO Neurons

Model LSO neurons were implemented using the Brian 2 Neural Simulator (Stimberg et al., 2019) in Anaconda Python 3.7 64-bit http://www.anaconda.com.

Code is available: https://github.com/AndrewBrughera/LSO_Transposed_Tones

Data and analysis scripts: https://doi.org/10.6084/m9.figshare.12798035.v1

IPD_ENV_, the interaural phase difference with respect to the amplitude modulation cycle of the transposed-tone was fixed within each simulation, and ranged from −180 to +180° in increments of 15°. At each IPD_ENV_, a different one of 24 envelope starting phase pairs was applied for each model LSO neuron within a population of 10 or 24 neurons. The 25 stimulus repetitions, each yielding spike times from the model ANFs, were each applied separately. Across the six populations of model LSO neurons, the same sets of input spike times were applied, except that for the slow model LSO neurons, a 1-ms delay was added to the spike-times driving contralateral inhibitory inputs, improving sensitivity to ITD_ENV_. In modeling with up to 120 LSO neurons per carrier frequency, we considered the 5600 neurons each in the left and right LSO (Kulesza, 2007), in just under 10 octaves of frequency range in human hearing from 20 to 20,000 Hz, yielding 140 neurons in each LSO per quarter-octave in equal log-spacing, without a bias toward high-frequency neurons.

Modeling LSO neurons, we extend an existing Hodgkin-Huxley-type model LSO neuron (Hodgkin & Huxley, 1952; Rothman & Manis, 2003; Wang & Colburn, 2012) to six populations: slow, moderate, HL-quick, brisk, fast, and very fast. All model LSO neurons have membrane conductances (Table 1) and reversal potentials, representing leakage (L) channels, sodium (Na^+^) channels, high-threshold potassium (K_HT_) channels, and hyperpolarization-activated cyclic nucleotide (h) channels for non-specific cations (reversal potentials: *V_L_* = −65 mV; *V_Na_* = +55 mV; *V_K_* = −70 mV; *V_H_* = −43 mV). Brisk, fast, and very fast model neurons also include a conductance (Table 1) and reversal potential for K_LT_ channels (*V_K_* = −70 mV). Slow model neurons use a model membrane for VCN stellate cells (Type 1c) (Rothman & Manis, 2003); moderate model neurons have increased conductances for h channels and leakage channels; and HL-quick model neurons have relatively high conductances for Na^+^, K_HT_, h and leakage channels. Fast model neurons use a model membrane for VCN bushy cells (Type 2) (Rothman & Manis, 2003), previously applied to LSO modeling (Wang & Colburn, 2012). The brisk model LSO neuron has half the K_LT_ and h conductances of the fast model neuron. The very fast model LSO neuron has double the Na^+^, K_HT_, K_LT_, and h conductances of the fast model neuron. Maximum conductance values (Table 1) were not adjusted for temperature. Voltage-sensitive gate activation and inactivation equations for ion channels (Na^+^, K_HT_, K_LT_, h) are identical to those in Rothman and Manis (2003c) including time constants divided by the Q_10_ temperature factor of 3^(*T* – 22)/10^; *T* is equal to human body temperature, 37°C. Consistent across model neurons are the threshold for counting action potentials (*V_AP-THRESHOLD_* = −30 mV), and the reversal potentials of excitatory synapses (*V_E_* = 0 mV) and inhibitory synapses (*V_I_* = −90 mV). In faster model neurons, synapses are made faster and stronger, as inhibitory synapses maintain 10 times the strength, and 2 times the duration, of excitatory synapses (Table 1). An input spike from a model ANF to an excitatory synapse increments, by the excitatory synaptic strength Δ*g_E_*, the total inhibitory synaptic conductance (*g_E_*) which decays exponentially with time constant *τ_E_* while producing excitatory synaptic current *i_E_* = *g_E_*(*V_E_* - *v_M_*), where *v_M_* is the membrane potential. Similarly, an input spike from a model ANF to an inhibitory synapse increments, by the inhibitory synaptic strength Δ*g_I_*, the total inhibitory synaptic conductance (*g_I_*) which decays exponentially with time constant *τ*_I_ while producing inhibitory synaptic current *i_I_* = *g_I_(V_I_* - *v_M_*), with the reversal potential *V_I_* below resting potential (*V_REST_*) to send current out of the model neuron.

**Table 1.** Parameters varying across model LSO neurons: maximum conductances (*g_max_*) for ion channels, membrane capacitance, synaptic strengths, and synaptic time constants

|  | Model LSO neurons: |  |  |  |  |  |
| --- | --- | --- | --- | --- | --- | --- |
| Ion-channel conductances: | Slow | Moderate | HL-Quick | Brisk | Fast | Very Fast |
| $g_{max_{Na}}$ | 1000 | 1000 | 2000 | 1000 | 1000 | 2000 |
| $g_{max_{KHT}}$ | 150 | 150 | 300 | 150 | 150 | 300 |
| $g_{max_{KLT}}$ | 0 | 0 | 0 | 100 | 200 | 400 |
| $g_{max_h}$ | 0.5 | 1.85 | 24 | 10 | 20 | 40 |
| $g_{Leak}$ | 2 | 7.4 | 24 | 2 | 2 | 2 |
| Membrane capacitance, $C_M$ | 12 | 21.7 | 12 | 12 | 12 | 12 |
| Excitatory synapses: |  |  |  |  |  |  |
| Strength, $\Delta g_E$ (nS) | 0.7 | 4.5 | 5 | 3 | 5 | 10 |
| Time Constant, $\tau_E$ (ms) | 1.5 | 1.5 | 0.5 | 1 | 0.5 | 0.2 |
| Inhibitory synapses: |  |  |  |  |  |  |
| Strength, $\Delta g_I$ (nS) | 7 | 45 | 50 | 30 | 50 | 100 |
| Time Constant, $\tau_I$ (ms) | 3 | 3 | 1 | 2 | 1 | 0.4 |

### Membrane impedance as a function of frequency in model LSO neurons

We calculated subthreshold membrane impedance as functions of frequency in each type of model LSO neuron (slow, moderate, HL-quick, brisk, fast, and very fast): based on the membrane potential resulting from an injected transmembrane current (i_Zap_), a “zap” stimulus containing a linear frequency sweep from 1 to 2000 Hz, beginning at time *t* = 0 in the equation below, with a duration *d* of one second (Table 2) (Hutcheon & Yarom, 2000; Puil et al., 1986; Remme et al., 2014).

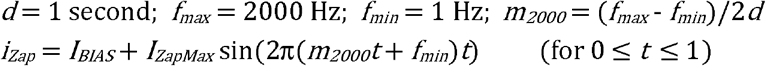

**Table 2.**
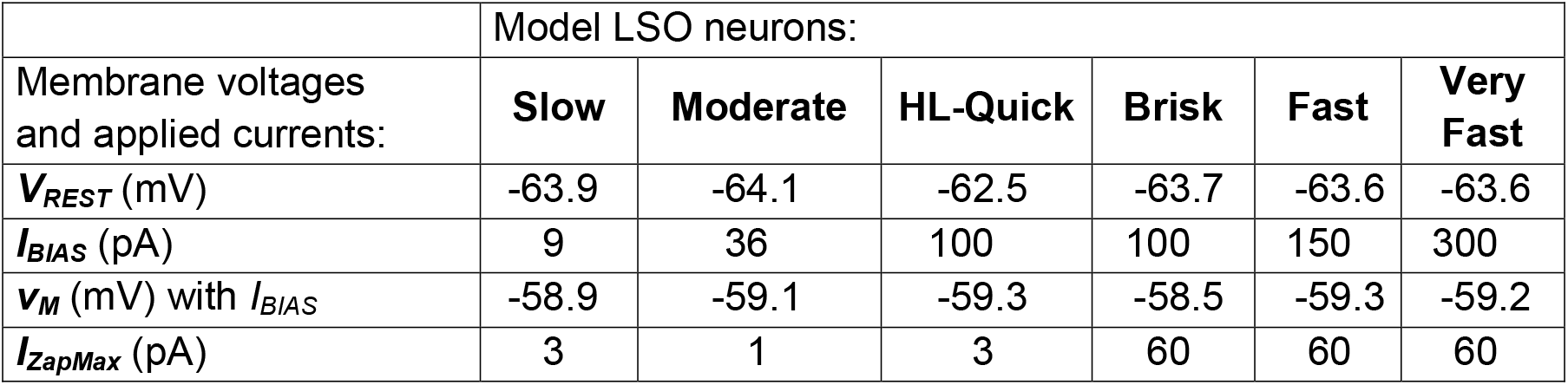
Current and voltage during membrane impedance determination

|  | Model LSO neurons: |  |  |  |  |  |
| --- | --- | --- | --- | --- | --- | --- |
| Membrane voltages and applied currents: | Slow | Moderate | HL-Quick | Brisk | Fast | Very Fast |
| $V_{REST}$ (mV) | -63.9 | -64.1 | -62.5 | -63.7 | -63.6 | -63.6 |
| $I_{BIAS}$ (pA) | 9 | 36 | 100 | 100 | 150 | 300 |
| $v_M$ (mV) with $I_{BIAS}$ | -58.9 | -59.1 | -59.3 | -58.5 | -59.3 | -59.2 |
| $I_{ZapMax}$ (pA) | 3 | 1 | 3 | 60 | 60 | 60 |

Prior to the “zap” stimulus, the model membrane settled for 50 ms to its resting potential near −64 mV. Then an inward bias current (*I_BIAS_*) was applied with magnitude such that *v_M_* increased to near −59 mV. This bias current was applied alone for 450 ms, and then maintained as the “zap” frequency sweep in membrane current (*i_ZAP_*) was also applied. For the HL-quick model neuron, we also examined the effect of reduced duration in bias current before *i_ZAP_*. Membrane impedance as a function of frequency was calculated using the fast Fourier transform (FFT). FFTs of the injected transmembrane current, and the resulting membrane voltage, were computed over the duration of the frequency sweep. The FFT of voltage was divided by the FFT of current, yielding the FFT of membrane impedance. The magnitude of this FFT was plotted, showing the magnitude of membrane impedance as a function of frequency.

### Neural Just-Noticeable Differences (JNDs) in Interaural Time Difference (ITD)

We calculated neural JNDs from Fisher information in the spike rates of our model LSO neurons (shown in Figs. 3 and 4) analyzed as functions of ITD_ENV_. For each of 25 stimulus presentations of a 300-ms transposed tone, we calculated a JND in ITD_ENV_ for each population of 24 model LSO neurons, based on a method of neuronal JND(IID) calculation (Brown & Tollin, 2016), which we extended to a neuronal population:

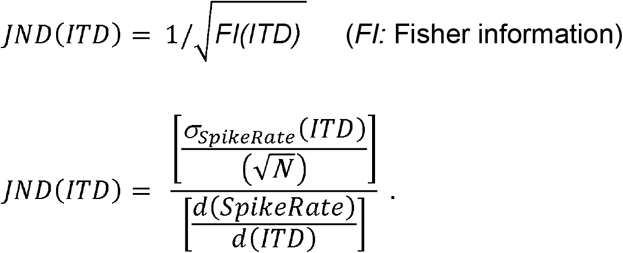

**Fig. 3.**
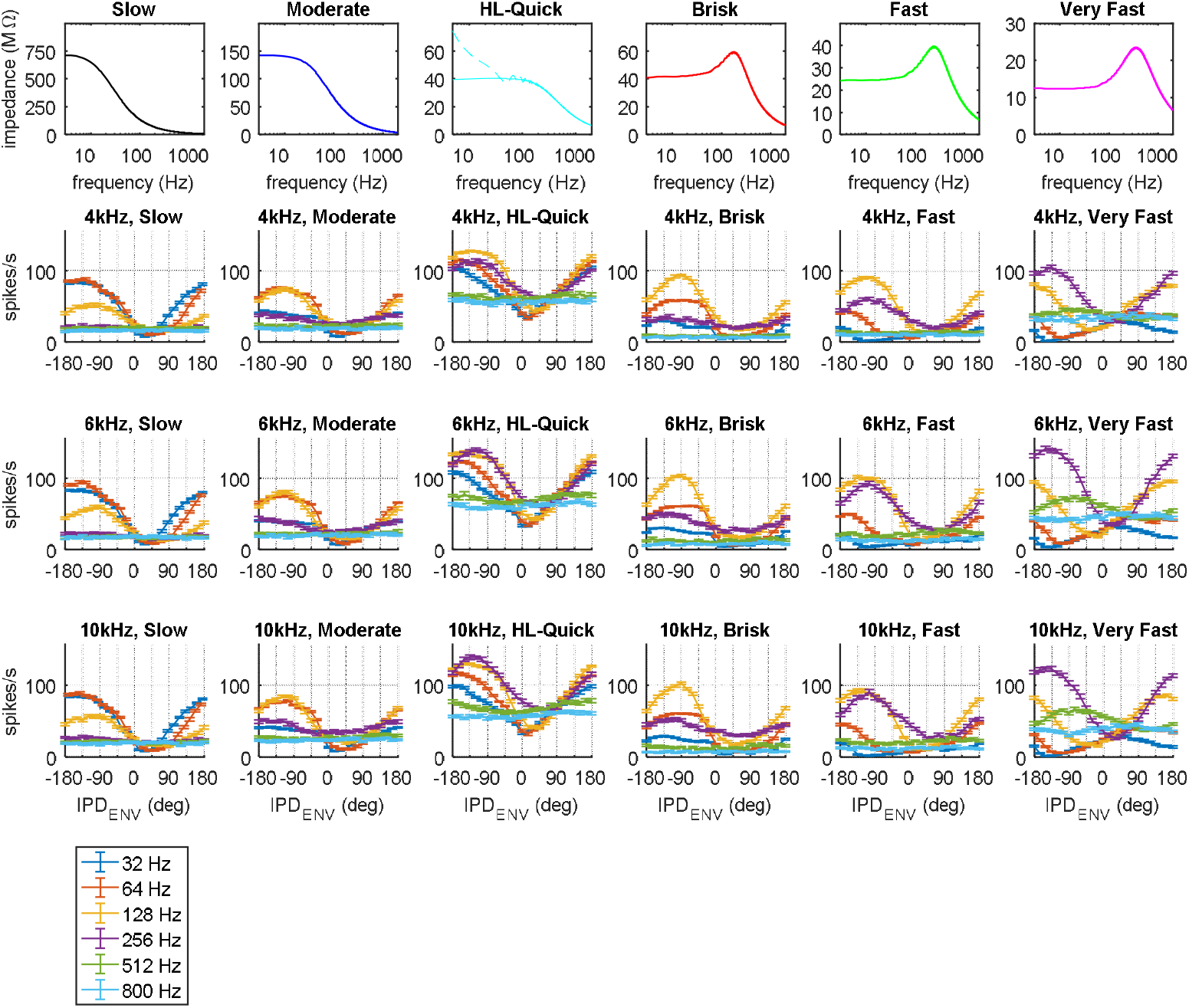
Membrane impedance and responses to transposed tones in model LSO neurons. Top row: Membrane impedance magnitude as functions of frequency in model LSO neurons of different membrane time constants: slow, moderate, HL-quick, brisk, fast, and very fast. Slow, moderate and HL-quick model neurons have non-resonant, low-pass membrane impedances. Brisk, fast, and very fast model neuronal membranes have model K_LT_ channels and resonant, low-pass membrane impedances. Lower 3 rows: Responses to on-frequency listening in populations of 24 model LSO neurons of each membrane speed: mean spike rate and standard error of the mean as functions of IPD_ENV_ (interaural phase difference with respect to the amplitude-modulation cycle). The title of each panel gives the stimulus carrier frequency, and membrane speed of the model LSO neurons. For on-frequency listening, the characteristic frequency (CF, the frequency of lowest spike-threshold in SPL) of model ANFs driving the model LSO neurons, was set equal to the carrier frequency. Responses are to the 25^th^ stimulus repetition, a separate curve for each modulation rate. The legend within the figure shows modulation rates

**Fig. 4.**
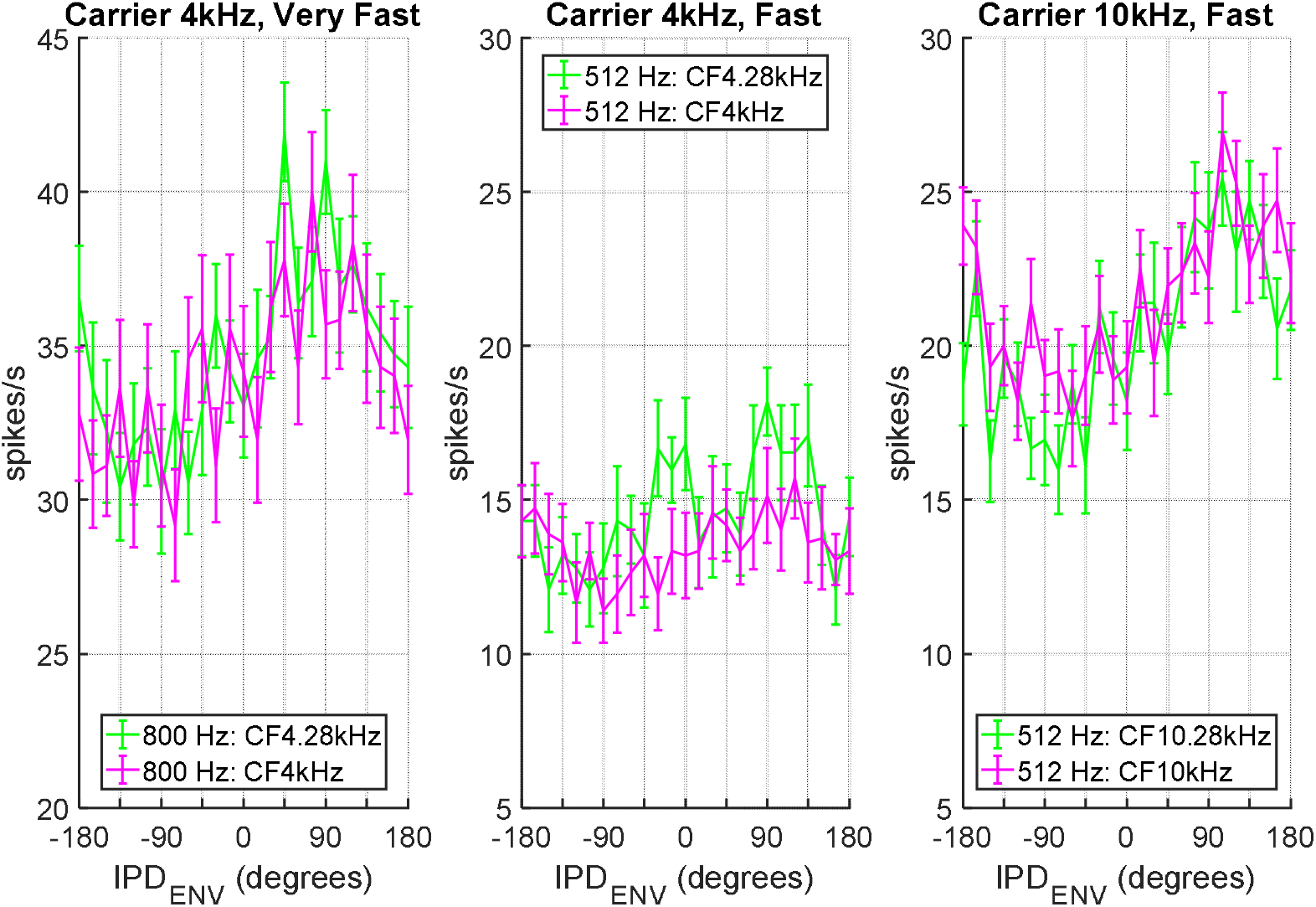
Model LSO neurons: on- and off-frequency listening. Mean spike rates as functions of IPD_ENV_ in populations of 24 model LSO neurons, for three stimulus-and-model conditions in which off-frequency listening produced a measurable JND and on-frequency listening did not (stimulus repetition 25, for all conditions). The title of each panel gives the carrier frequency of the transposed tone stimulus, and the membrane speed of model LSO neurons. The legend within each panel shows modulation rate and CF of the model neurons. Error bars show standard error of the mean spike rate. The criterion for a valid JND is agreement in the algebraic sign in the slope of spike rate with respect to IPD_ENV_ from −45° to 0°, for at least 19 of 25 stimulus repetitions. For these three conditions of off-frequency listening (CF between the carrier frequency and modulation sideband), as IPD_ENV_ increased from −45° to 0°, the changes in spike rate were positive, agreeing with the trend in slope across the conditions. For on-frequency listening (CF equal to the carrier frequency), as IPD_ENV_ increased from −45° to 0°, the changes in spike rate were negative, positive, and near zero for one condition each, suggesting how on-frequency listening failed to meet the criterion for a measurable JND within each condition

The arithmetic standard deviation in spike rate, σ_SpikeRate_, was calculated over the *N* (24 or 10) model neurons in the population. Thus the bracketed numerator is the standard error of the mean spike rate for this population, calculated as the root-mean-square of the standard errors of mean spike rate at IPD_ENV_ equal to −45° and 0°. The bracketed denominator is the derivative of mean spike rate with respect to ITD_ENV_, calculated as a linear slope over the interval of ITD_ENV_ equivalent to IPD_ENV_ from −45° to 0°. Hence, the JND is equal to the standard error of the mean spike rate, divided by the derivative of mean spike rate with respect to ITD_ENV_. Agreement in the algebraic sign of the derivative in at least 19 of the 25 stimulus repetitions is the criterion for valid JND statistics in each stimulus condition. If this criterion was not met, the JND was considered to be unmeasurably high (the task was impossible). If this criterion was met, the geometric mean and standard deviation of JND were then calculated from the absolute values of all finite values of JND from the 25 stimulus repetitions. Typically all 25 of 25 JND values were finite, and the minimum was 23 of 25.

## Results

### Membrane impedances and frequency responses of model neuronal membranes

We calculated subthreshold membrane impedance as functions of frequency for model LSO neurons of different speeds (slow, moderate, HL-quick, brisk, fast, and very fast; according to their membrane time constants, see below) (Fig. 3), based on the membrane potential evoked by injected transmembrane current containing a linear frequency sweep from 1 to 2000 Hz, over a duration of one second (Hutcheon & Yarom, 2000; Puil et al., 1986; Remme et al., 2014). The membrane impedance of each model neuron is a low-pass function of frequency: non-zero at frequency zero and ultimately decreasing with higher frequency, describing membrane voltage that responds to a static input current, and ultimately responds progressively less to input currents of higher frequency.

Our slow, moderate, and HL-quick model LSO neurons have non-resonant, low-pass membrane impedances. The slow model neuron has high membrane impedance, maximum magnitude 713 mega-ohms (MΩ) at 3 Hz, magnitude 704 MΩ at 1 Hz, and a low-pass cut-off (−3 dB) frequency of 21 Hz, obtained by applying a model membrane for a ventral-cochlear-nucleus (VCN) stellate cell (Type 1c) (Rothman & Manis, 2003). With higher membrane impedance, lower cut-off frequency, and slower membrane time constant than observed LSO neurons, the slow model neuron will illustrate that even the slowest of LSO neurons can be sensitive to ITD_ENV_. The moderate model neuron resembles typical non-resonant LSO neurons (Barnes-Davies et al., 2004; Remme et al., 2014), with moderate membrane impedance, maximum magnitude 143 MΩ at 3 Hz, magnitude 142 MΩ at 1 Hz, and a low-pass cut-off frequency of 53 Hz. The HL-quick model neuron resembles an observed medial LSO neuron (Remme et al., 2014), with low membrane impedance (Fig. 3, top row: solid cyan line), maximum magnitude 41 MΩ at 35 Hz, magnitude 37 MΩ at 1 Hz, and a low-pass cut-off frequency of 395 Hz. With reduced duration, 190 ms vs. 450 ms, in the bias current before adding the frequency sweep (see Methods), membrane impedance of the HL-quick model neuron increases below 60 Hz (Fig. 3, top row: dashed cyan line), consistent with the membrane impedance measured in the real neuron.

Our brisk, fast, and very fast model neurons feature model K_LT_ channels, and resonant low-pass membrane impedances, as in LSO neurons expressing K_LT_ channels (Barnes-Davies et al., 2004; Remme et al., 2014). K_LT_ channels produce transmembrane current that opposes depolarization (Manis & Marx, 1991), analogous to inductive current in an underdamped parallel RLC circuit, and producing a similar characteristic resonance peak in membrane impedance (Nilsson & Riedel, 2008). The fast model LSO neuron (Rothman & Manis, 2003; Wang & Colburn, 2012) has low membrane impedance, with peak magnitude 40 MΩ at its resonance frequency of 249 Hz. The brisk model LSO neuron has higher membrane impedance, with a less prominent peak in magnitude of 60 MΩ at a lower resonance frequency, 173 Hz. The very fast model LSO neuron has lower membrane impedance, with a more prominent peak in magnitude of 24 MΩ at a higher resonance frequency, 355 Hz.

Each type of model LSO neuron is distinct in the frequency response of its membrane impedance (Fig. 3), with the resulting calculated membrane time constant: slow (7.6 ms), moderate (3.0 ms), HL-quick (0.48 ms), brisk (0.92 ms), fast (0.64 ms), and very fast (0.45 ms). In non-resonant membranes, a first-order system, the calculated membrane time constant is equal to the reciprocal of the low-pass cut-off frequency (the −3-dB frequency, in radians per second); in resonant membranes, a second order system, the calculated membrane time constant is equal to the reciprocal of the resonance frequency (in radians per second) (Nilsson & Riedel, 2008). A neuron with a faster membrane time constant can effectively pass higher frequencies from its input current to its membrane potential. With faster membrane time constants and lower membrane impedance, model synapses were made faster and stronger to maintain action potentials (Sterenborg et al., 2010). Across neuronal speeds, model inhibitory synapses maintained double the time constant, and 10 times the strength, of model excitatory synapses.

### Membrane frequency response influences ITD_ENV_ sensitivity in model LSO neurons

For populations of 24 model LSO neurons of each speed (slow, moderate, HL-quick, brisk, fast, and very fast), binaurally stimulated by transposed tones, we plotted mean spike rate as a function of envelope IPD (IPD_ENV_), the interaural phase difference with respect to the amplitude-modulation cycle (Fig. 3). The transposed tones were of 300-ms duration, with maximum sound pressure level (SPL) 75 dB RMS, at carrier frequencies of 4, 6, and 10 kHz, and modulation rates of 32, 64, 128, 256, 512, and 800 Hz. The transposed tones stimulate a bilateral-auditory-periphery model containing stochastic model auditory-nerve (AN) fibers (Zilany et al., 2014) of characteristic frequency (CF, the frequency of lowest spike-threshold in SPL) equal to the stimulus carrier frequency. Model ANFs directly drive the 40 ipsilateral excitatory synapses and 8 contralateral inhibitory synapses of each model LSO neuron (Fig. 1B), where membrane impedance low-pass filters the inputs driven by amplitude-modulated sound, limiting the upper modulation rate for sensitivity to ITD_ENV_.

Reasonably consistent with the frequency response of its membrane impedance, each model population of LSO neurons generates spike rates that are most sensitive to IPD_ENV_ for a particular range of modulation rate. Slow model neurons are most sensitive for modulation at 32 and 64 Hz. Moderate model neurons are most sensitive for modulation at 64 and 128 Hz, with lower sensitivity at 32 Hz and still less sensitivity at 256 Hz. HL-quick model neurons are most sensitive for modulation at 128 and 256 Hz, with lower sensitivity for 32-Hz and 64-Hz modulation. At 32-Hz modulation, slow model neurons have steep spike-rate functions of IPD_ENV_ (rate-IPD_ENV_ functions), spiking from near zero to three times during the 31.25-ms modulation cycle; moderate model neurons do not spike repeatedly, yet their rate-IPD_ENV_ functions approach similar steepness near zero IPD_ENV_. As modulation rate increases to 512 and 800 Hz, slow, moderate, and HL-quick model neurons become quite insensitive to IPD_ENV_. Brisk model neurons are most sensitive for modulation at 64 and 128 Hz, with lower sensitivity at 32 and 256 Hz. Fast model neurons are most sensitive for modulation at 128 and 256 Hz. Very fast model neurons are most sensitive for modulation at 256 Hz, and sensitive at 128 and 512 Hz. With decreasing modulation rates below 128 Hz, the rate-IPD_ENV_ functions of fast and very fast model neurons decrease in slope, and their steepest slopes occur far from zero IPD_ENV_.

At high modulation rates of 512 and 800 Hz, modulation output is increased by band-pass filters centered above the carrier frequency (Monaghan et al., 2015). We investigated this ‘off-frequency listening’ at 512-Hz and 800-Hz modulation, using populations of brisk, HL-quick, fast, and very fast model LSO neurons, stimulated by model ANFs with CF between the carrier frequency and the first upper modulation sideband of each transposed tone. At these high modulation rates, for brisk model neurons, neither this off-frequency listening, nor the original on-frequency listening (CF at the carrier frequency), produced measurable JNDs in ITD_ENV_. For HL-quick model neurons, off-frequency listening did not produce a measureable JND, and on-frequency listening produced a measurable JND in only one condition. For fast and very fast model neurons, compared with model neurons of the same membrane speed with CFs at the carrier frequency, off-frequency listening produced some small increases in sensitivity to ITD_ENV_ (Fig. 4), sufficient generally to improve neural JNDs in ITD_ENV_, including three conditions requiring off-frequency listening to produce a measurable JND (JNDs shown below in Fig. 5). For fast and very fast model neurons, each condition in which on-frequency listening produced a measurable JND, off-frequency listening also produced a measurable JND.

**Fig. 5.**
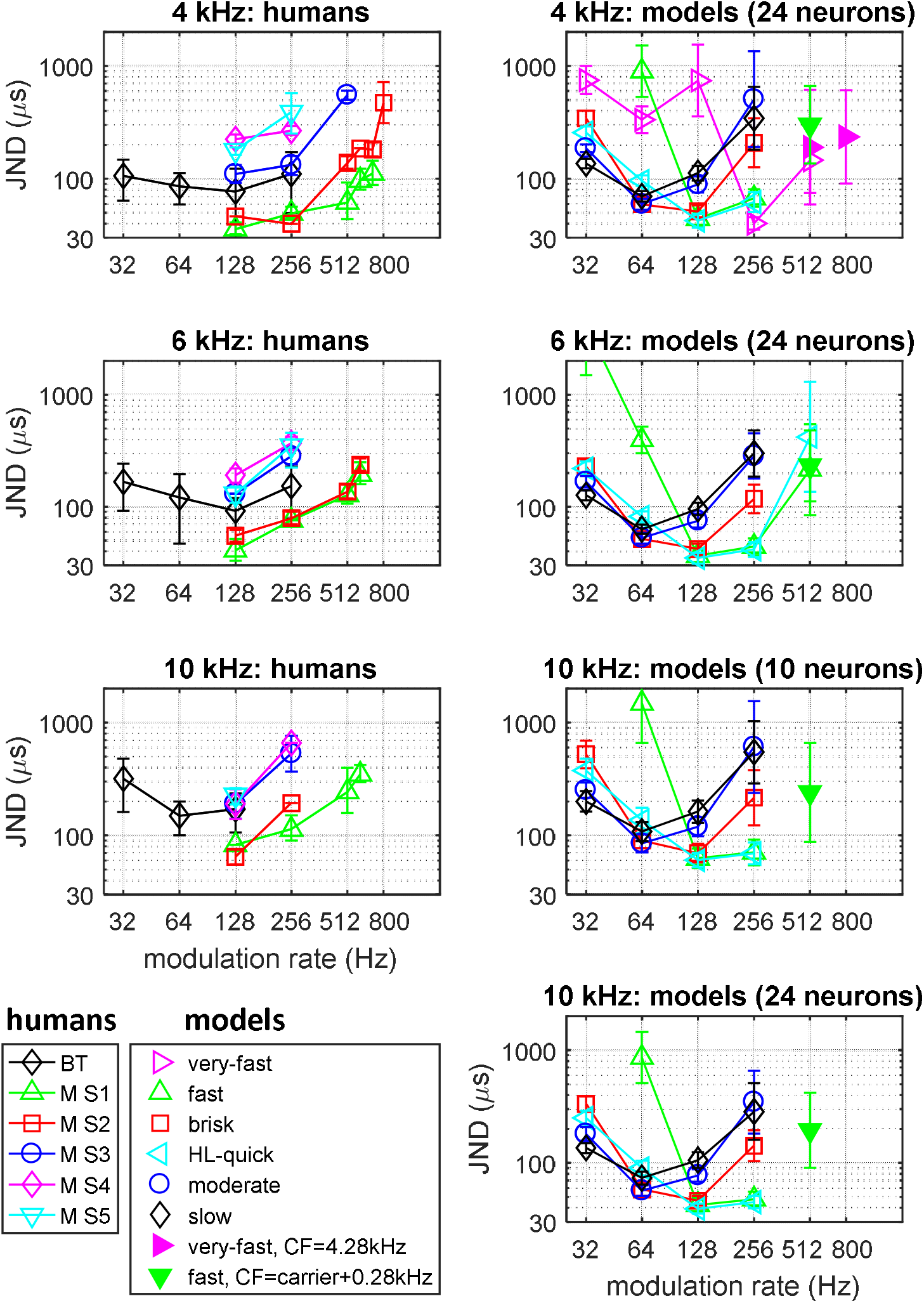
Just-noticeable differences (JNDs) in ITD_ENV_ as functions of modulation rate in transposed tones at carrier frequencies of 4, 6, and 10 kHz. The title of each panel includes the carrier frequency. Left column: Human ITD_ENV_ discrimination thresholds replotted from Bernstein and Trahiotis (2002) (BT, arithmetic mean and standard deviation across listeners) and from (Monaghan et al., 2015) (geometric mean and standard deviation for each listener, M S1 thru M S5). Right column: Neural JNDs in ITD_ENV_ calculated for populations of 10 or 24 model LSO neurons (geometric mean and standard deviation across 25 stimulus repetitions). These populations are distinct in membrane time constant: slow, moderate, HL-Quick, brisk, fast, and very fast. Model neurons generally applied on-frequency listening: CF equal to the carrier frequency. At high modulation rates, additional populations of fast and very fast model neurons applied off-frequency listening: CF between the carrier frequency and modulation sideband, as shown in the legend within the figure

Overall, neural tuning for ITD_ENV_ depends on membrane speed. Model neurons with relatively slow membranes—characterized by high input impedances and low-pass electrical filters of low cut-off frequency—are most suited to extracting ITDs from slowly modulated sounds, and neurons with faster membranes less so. As modulation rate increases, model neurons with faster membrane properties—low input impedance with a high-frequency resonance or cut-off—show relatively greater sensitivity to ITD_ENV_ compared to neurons with slow membrane properties.

### Just-noticeable differences in ITD_ENV_: humans and model LSO neurons

Using the spike-rate responses to transposed tones, we calculated JNDs in ITD_ENV_ for our model LSO populations, and compared them with human ITD_ENV_ discrimination thresholds from published reports (Fig. 5). For the models, modulation rates were 32, 64, 128, 256, 512, and 800 Hz. In the human studies, modulation rates ranged from 32 to 512 Hz (Bernstein & Trahiotis, 2002), and from 128 to 800 Hz (Monaghan et al., 2015).

For each stimulus repetition and model population, a neural JND (Brown & Tollin, 2016) was calculated as the standard error of the mean spike rate, divided by the derivative of mean spike rate with respect to ITD. Agreement in the algebraic sign of the derivative in at least 19 of the 25 stimulus repetitions was required. If this criterion was met, the geometric mean and standard deviation of the absolute values in JND were then calculated and plotted, using all finite JNDs from the 25 stimulus repetitions. If this criterion was not met, the JND was considered to be unmeasurably high (i.e. the task was impossible).

For transposed tones at 4 kHz, human JNDs in ITD_ENV_ are lowest (performance is best) at 128-Hz modulation. At higher modulation-rates, performance worsens (JNDs increase), and eventually the task becomes impossible: all the human listeners were able to perform the task at 256-Hz modulation; 5 of 9 listeners were capable at 512-Hz modulation; 2 of 5 at 600-Hz and 700-Hz; and 1 of 5 at 800-Hz (Bernstein & Trahiotis, 2002; Monaghan et al., 2015). Slow and moderate model LSO neurons generally reflected a competent level in average human performance for modulation from 32 to 128 Hz (Bernstein & Trahiotis, 2002), but then worsened to reflect below-average human performance at 256-Hz modulation. At modulation rates above 32 Hz, top human performance was reflected by the best-performing model populations: this included brisk and moderate model neurons at 64-Hz modulation, HL-quick and fast model neurons at 128-Hz modulation, and very fast model neurons at 256- to 800-Hz modulation, requiring off-frequency listening at 800-Hz. At 4 kHz, 256-Hz modulation is a condition where the range of model neuronal membranes produces JNDs spanning the largest human range of JNDs. At 128-Hz modulation, the condition of lowest human JND (Monaghan et al., 2015), model membrane speed does not account for poor human performance, as very fast model neurons performed still worse, and slow-to-fast model populations performed better.

For transposed tones at 6 kHz, human JNDs in ITD_ENV_ are again lowest at 128-Hz modulation, and increase with higher modulation rates. Human JNDs increase slightly overall compared with 4 kHz, yet all human listeners capably performed the task at 256-Hz modulation; 4 of 9 were capable at 512-Hz modulation; and 2 of 5 at 600-Hz (Bernstein & Trahiotis, 2002; Monaghan et al., 2015). Consistent with broadening auditory filters at higher CF better passing the spectral modulation sidebands of amplitude-modulated sounds, JNDs of the model populations with 24 neurons decreased slightly, compared with those for transposed tones at 4 kHz. Slow and moderate model neurons reflected average to above-average human performance from 32- to 128-Hz modulation, and below-average performance at 256-Hz modulation. At modulation rates above 32 Hz, the best performing model populations generally reflected top human performance: which was approached by moderate and brisk model neurons at 64-Hz modulation, reflected by HL-quick, brisk and fast model neurons at 128-Hz modulation, spanned by brisk to fast and HL-quick model neurons at 256-Hz modulation, and matched by fast model neurons at 512-Hz modulation using either on-frequency or off-frequency listening. At 128-Hz modulation, a condition with generally very low JNDs for humans (Bernstein & Trahiotis, 2002; Monaghan et al., 2015), model membrane speed does not account for poor human performance, as slow to fast model populations all performed better. Consistent with the lower proportion of fast resonant LSO neurons at higher CFs (Remme et al., 2014): slow to fast model neurons were sufficient to match human ITD_ENV_ sensitivity; at 6 and 10 kHz, very fast model neurons were not required to match human performance.

For transposed tones at 10 kHz, human JNDs increase relative to those at 6 kHz. For humans and model neurons, JNDs in ITD_ENV_ at 10 kHz are generally lowest at 64-Hz and 128-Hz modulation. At higher modulation rates, performance worsens: 7 of 9 human listeners capably performed the task at 256-Hz modulation; 2 of 9 were capable at 512-Hz modulation; and 1 of 5 at 600-Hz (Bernstein & Trahiotis, 2002; Monaghan et al., 2015). Maintaining populations of 24 model neurons, the top-performing model neurons generally exceeded top human performance. Reducing each population to 10 model neurons restored the match of human performance. Slow and moderate model neurons reflected average to above-average human performance from 32- to 128-Hz modulation, and below-average performance at 256-Hz modulation. Top human performance was approached by moderate and brisk model neurons at 64-Hz modulation; reflected by HL-quick, fast, and brisk model neurons at 128-Hz modulation; spanned by brisk to fast and HL-quick model neurons at 256-Hz modulation; and matched by fast model neurons at 512-Hz modulation using off-frequency listening. At 128-Hz and 256-Hz modulation, model membrane speed largely accounts for the range of human performance. However, at 64-Hz modulation, which has the lowest measured mean JND at 10 kHz for humans (Bernstein & Trahiotis, 2002), model membrane speed does not account for poor human performance, as fast model neurons performed still worse, and slow, moderate, HL-quick, and brisk model populations performed better. Considering 4-, 6-, and 10-kHz carriers, at modulation rates where human ITD_ENV_ sensitivity is generally the best, model membrane speed does not account for poor human performance; then at higher modulation rates where human ITD_ENV_ sensitivity decreases, the range in model membrane speeds produces JNDs in ITD_ENV_ which reflect the range of human performance.

Consistent with the paradoxical reduction in ITD_ENV_ sensitivity with increasing sound frequency observed behaviorally in humans (Bernstein & Trahiotis, 2002; Monaghan et al., 2015): reducing each population to 10 model neurons at 10 kHz increased JNDs relative to those at 6 kHz for the same modulation rate, and provided a similar fit to human performance to those of the populations of 24 model neurons at 4 and 6 kHz.

#### Effects of carrier frequency

As carrier frequency increases, sensitivity to ITD_ENV_ in human listeners decreases (JNDs increase). This decreasing sensitivity with increasing sound frequency is paradoxical, as cochlear band-pass filters broaden in absolute bandwidth with higher CF (Glasberg & Moore, 1990), thereby more strongly passing the modulation sidebands of amplitude-modulated stimuli. Consistent with increasing peripheral bandwidth, at 256-Hz modulation with increasing carrier frequency from 4 to 6 kHz, for brisk and fast model neurons we find slightly elevated modulation in spike rate as a function of IPD_ENV_ (Fig. 3), and slightly reduced (improved) JNDs in ITD_ENV_ (Fig. 5). Similar increases in spike-rate modulation occur in very fast model neurons for 800-Hz modulation, for increasing carrier frequencies from 4 to 6 to 10 kHz (Fig. 3). For slow to fast model neurons, the steady JNDs between 6 and 10 kHz when maintaining 24 model neurons per population (Fig. 5) suggest that the model cochlear filters have sufficient bandwidth at 6 kHz to pass relevant sidebands for modulation up to 256 Hz. The increasing human JNDs (decreasing sensitivity) with higher carrier frequency may be explained by decreasing numbers of stimulated auditory neurons with higher CF, as suggested to occur in ”hidden” hearing loss (Plack et al., 2014). When the number of model neurons was reduced to 10 per population at 10 kHz (maintaining 24 model neurons per population at 4 and 6 kHz), JNDs increased from 6 to 10 kHz (Fig. 5), reproducing the paradoxical reduction in ITD_ENV_ sensitivity with increasing sound frequency.

#### Effects of modulation rate

For humans and our models, JNDs in ITD_ENV_ are lowest (performance is best) at 64-Hz, 128-Hz, or 256-Hz modulation. Halving the modulation rate, the same change in spike rate for IPD_ENV_ between −45° and 0° (Fig. 3) produces double the JND in ITD_ENV_ (Fig. 5). With increasing modulation rate, although this effect becomes an advantage, it is overcome by decreasing membrane impedance that reduces the response of a model neuronal membrane to synaptic currents: performance worsens, and eventually the task becomes impossible. At modulation rates of 256 Hz and above, the large variance in human ITD_ENV_ sensitivity including unmeasurable JNDs is reflected by the JND range of the model neuronal populations.

#### Effects of off-frequency listening

Modulation at 512 and 800 Hz in the output of auditory-like gammatone filters is greater for filters centered above the carrier frequency, compared with gammatone filters centers at the carrier (Monaghan et al., 2015). Accordingly, with off-frequency listening, modeled by applying inputs from model ANFs of CF between the carrier frequency and first upper modulation sideband: at 512-Hz and 800-Hz modulation, JNDs in ITD_ENV_ are generally lower (better performance) for fast and very fast model LSO neurons, when compared with JNDs for model neurons of the same speed with inputs of CF at the carrier frequency. For multiple stimulus conditions, off-frequency listening provided a measurable JND, when on-frequency listening did not (Fig. 5). This suggests that the very best listeners, in terms of ITD_ENV_ discrimination performance, can access relatively small fluctuations in neural activity in frequency channels beyond that of the stimulus center-frequency to perform discrimination tasks, and that this off-frequency listening performance still requires the existence of fast and very-fast LSO neurons to extend to frequencies beyond the range of average-performing listeners.

## Discussion

We explored sensitivity to ITD_ENV_ in computationally modeled LSO neurons in potentially explaining human behavioral discrimination of ITD_ENV_ in high-frequency amplitude-modulated sound. In particular, we evaluated the influence of electrical impedance of neuronal membranes on discrimination performance as a function of the modulation rate and carrier frequency of sound. Transposed tones stimulated an auditory-periphery model bilaterally innervating the model LSO neurons. Collectively reflecting human abilities to discriminate ITD_ENV_, neural JNDs in ITD_ENV_ were calculated for model neuronal populations, distinct in membrane frequency response reflecting the LSO range, with time constants from slow to very fast. Within these model neurons, electrical membrane impedances similar to those in LSO neurons decrease with increasing frequency, and limit the upper modulation rate for sensitivity to ITD_ENV_, by low-pass filtering binaural inputs driven by amplitude-modulated sound. Yet, rather than very fast model neurons with the widest bandwidth being best overall, each speed of model neuron produced the lowest JNDs in ITD_ENV_ for a particular range of modulation rate: slow model neurons at 32 Hz; moderate model neurons at 64 Hz; brisk model neurons from 64 to 128 Hz; fast and HL-quick model neurons from 128 to 256 Hz; and very fast model neurons from 256 to 800 Hz. Across modulation rates, top sensitivity to ITD_ENV_ leveraged model neurons of all speeds from slow to very fast. Although the slowest LSO neurons may not be quite as slow as the slow model neurons, which illustrate the capacity for ITD_ENV_ sensitivity in slow neurons, JNDs at 32-Hz modulation were nearly as low for moderate model neurons, suggesting that moderate LSO neurons may combine with slightly slower LSO neurons to lower the JND at 32-Hz modulation. With compatible inputs and synapses, our models suggest that ITD_ENV_ can be well encoded collectively by LSO neurons of the observed range in membrane properties.

Our model neuronal populations reflect the dependence of human ITD_ENV_ sensitivity on the carrier frequency and modulation rate of sound, including the ultimate reduction in sensitivity with increasing modulation rate, and the paradoxical reduction in sensitivity with increasing carrier frequency, modeled here by the combination of reduced top membrane speed, and a reduced number of stimulated neurons as suggested to occur in ”hidden” hearing loss (Plack et al., 2014). In each stimulus condition, across carrier frequency (4-10 kHz) and modulation rate (32-800 Hz), top-range human sensitivity to ITD_ENV_ is generally reflected by the top-performing population of model LSO neurons. Off-frequency listening in fast and very fast model neurons, with CFs between the carrier frequency and modulation sideband, helped to extend ITD_ENV_ sensitivity to modulation rates above 500 Hz. Reflecting the range in low-pass-filter properties of LSO neuronal membranes (Remme et al., 2014), the model neuronal populations collectively reproduce the largest (ten-fold) variation in measurable JNDs for ITD_ENV_ across human listeners, which occurs at 4 kHz with 256-Hz modulation (Monaghan et al., 2015), and is larger than the variation in behavioral thresholds for ITD_TFS_ (Bernstein & Trahiotis, 2002).

Early interpretation of differences in human ability to exploit ITD_TFS_ and ITD_ENV_ focused on differences in the peripheral representation of low- and high-frequency sounds. The Colburn and Esquissaud (1976) hypothesis posits equivalent sensitivity to ITD_TFS_ and ITD_ENV_, if peripheral representations of low-frequency sounds and high-frequency modulated sounds are matched prior to binaural integration. Bernstein and Trahiotis (2002) explored this concept employing transposed tones of modulation frequencies equal to the frequencies of low-frequency tones. Despite asserting approximately equivalent sensitivity for ITD_TFS_ and ITD_ENV_, human discrimination thresholds in ITD_ENV_ were actually substantially better at the lowest modulation frequencies for transposed tones (32 and 64 Hz) than they were for pure tones of 32 and 64 Hz (Bernstein & Trahiotis, 2002). A plausible explanation is that the relatively fast MSO neurons encoding ITD_TFS_ are less able to encode ITD information at 32 and 64 Hz, than LSO neurons with moderate to reasonably slow, more integrative, membrane properties when operating at similarly slow modulation rates.

Contrasting with improving performance in ITD_TFS_ up to at least 700 Hz and sensitivity to 1400 Hz (Brughera et al., 2013), as modulation frequencies increase beyond 128 Hz, Bernstein and Trahiotis (2002) suggested that a low-pass modulation filter, initially proposed with a corner frequency of 150 Hz, would explain the declining sensitivity to ITD_ENV_. While explaining average performance of their four listeners, this filter poorly fit individual listeners: one listener had comparable thresholds for ITD_ENV_ at 512-Hz modulation, and ITD_TFS_ thresholds at the same pure-tone frequencies; the remaining three listeners showed ten-fold increases in their ITD_ENV_ thresholds, or were unable to perform the task. Error bars, for the average performance of these four listeners, indicate their range of performance at 128 and 256 Hz is similar to that of individual thresholds for the five listeners in Monaghan et al. (2015). We modelled this decline in performance between listeners as decreasing access to LSO neurons with relatively high membrane speeds, which are produced *in vivo* by the expression of ion channels. Decreasing access to LSO neurons with high membrane speeds also helped reproduce the worsening of ITD_ENV_ performance with increasing CF. Considered across listeners and CFs, availability of LSO neurons with high membrane speeds would explain the fact that listeners with the lowest ITD_ENV_ thresholds at 128 and 256 Hz for a 4-kHz carrier frequency were invariably those for whom ITD_ENV_ remained discriminable at higher modulation rates and at higher carrier frequencies. The practical low-pass modulation filter is carrier-frequency dependent (Bernstein & Trahiotis, 2014), and varies across human listeners for the same modulation and carrier frequencies, consistent with our modelling data. This suggests a fundamental determinant of ITD_ENV_ sensitivity might be the ability of different listeners to access fast LSO membrane properties. Heterogeneity in the expression of ion channels is a systemic feature of physiology (Schulz et al., 2006; Veerman et al., 2017), suggesting a cogent explanation for the inter-subject variability.

### Complimentary cues in auditory spatial encoding

Categorical, cyclic, and non-cyclic continuous-scale encoding of auditory spatial cues may together support accurate sound-source localization. The brain may disambiguate the cyclic encoding of ITDs in the MSO and LSO (Batra et al., 1997), by applying non-cyclic encoding of IIDs by non-principal LSO neurons (Boudreau & Tsuchitani, 1967), and categorical encoding of onset IID by principal LSO neurons (Franken et al., 2018), combined with processing of categorical front/back and up/down spectral distinctions in pinnae acoustics (Langendijk & Bronkhorst, 2002). Consistent with recent definitions (Franken et al., 2018): principal LSO neurons refer to resonant LSO neurons with bipolar dendrites, and non-principal LSO neurons include non-resonant LSO neurons. Although suggesting a mechanism for observed psychophysical lateralization based on the leading ear when presented with confounding ITD cues (Stern & Shear, 1996; Thompson et al., 2006; von Kriegstein et al., 2008), onset-IID responses to unmodulated high-frequency tones in six principal LSO neurons (Franken et al., 2018) do not preclude the well-documented sensitivity of LSO neurons to ITD_ENV_ in high-frequency amplitude-modulated sound (Batra et al., 1997; Joris & Yin, 1995, 1998). Reflecting behavioral sensitivity, neural sensitivity to static ITD_ENV_ decreases with increasing modulation rate, yet can show clear modulation in spike rate at 500-Hz modulation, persisting at 750-Hz modulation (Joris, 1996). Depending on their inputs and membranes, principal or non-principal neurons may contribute to these responses.

### Stimuli, neurons, and encoding mechanisms differ between ITD_TFS_ and ITD_ENV_

Overall, MSO neurons encoding ITD_TFS_ have advantages over LSO neurons encoding ITD_ENV_. Temporally-precise MSO neurons tuned to low sound frequency encode ITD_TFS_ (Franken et al., 2015; Yin & Chan, 1990), applying binaural-coincidence-detection mechanisms sharpened by K_LT_ channels (Mathews et al., 2010), as dendritic mechanisms emphasize binaural relative to monaural coincidences (Dasika et al., 2007; Scott et al., 2010). With high-frequency amplitude-modulated sound, modulation rate is limited by the bandwidth of cochlear filters, and phaselocking in the AN is generally lower even for transposed tones than for low-frequency tones (Dreyer & Delgutte, 2006). Despite these peripheral effects, a high-frequency MSO neuron was quite sensitive to ITD_ENV_ for 1000-Hz modulation (Yin & Chan, 1990). Although some LSO neurons express K_LT_ channels (Barnes-Davies et al., 2004), LSO neurons do not benefit from binaural-coincidence detection, instead relying on the anti-coincidence of ipsilateral excitation and contralateral inhibition (Batra et al., 1997). LSO neurons with slowly-responding membranes lacking K_LT_ channels become increasingly common in higher-frequency LSO regions (Barnes-Davies et al., 2004; Remme et al., 2014). The temporal and frequency limits of LSO neurons appear to shape the modulation-rate limits of ITD_ENV_ sensitivity.

### Implications for listeners with bilateral cochlear implants (CIs, bCIs)

Unlike in normal-hearing listeners leveraging the advantages of ITD_TFS_, in bCI listeners, speech unmasking can be about equal for envelope cues, compared with fine-structure cues (van Hoesel et al., 2008). For bCI stimulation at high pulse rates typical of clinical CI processors, bCI listeners achieve about 9 dB of binaural unmasking when the temporal envelope is presented without fine structure using constant-rate pulses (Todd et al., 2019). This benefit from binaural envelope cues may be largely from IID with only minor support from ITD_ENV_, consistent with the much heavier weighting of IID than of ITD for localizing high-frequency sound, by the large majority of normal-hearing listeners (Macpherson & Middlebrooks, 2002).

Considerable effort is showing how bCI listeners might further benefit from binaural localization cues (Kan et al., 2019, 2015; Laback et al., 2011; Monaghan & Seeber, 2016; Stakhovskaya & Goupell, 2017), and notably in conveying electrical ITD to bCI listeners using bilateral pulse-burst stimulation (Srinivasan et al., 2020, 2018), which might be incorporated in conveying acoustic ITD_TFS_, ITD_ENV_, IID, and speech to bCI listeners. CI-stimulation rates up to 1000 or 1200 pulses per second would be sufficiently slow to convey coherent ITD cues using pulse bursts, and sufficiently fast to provide speech information (Fu & Shannon, 2000; Shader et al., 2020; Shannon et al., 2011). A bCI processor can be developed to present electrical pulses synchronous with the TFS of sound at each ear, bilaterally coordinated with pulse bursts, temporal interleaving across electrodes for low- to high-frequency sound, and refractory pauses for high-frequency channels.

## Declarations

### Ethics approval

For this strictly computational study, no approval was required.

### Consent to participate

For this strictly computational study, no consent was required.

### Consent for publication

For this strictly computational study, there are no human subjects from whom consent is required. All authors and responsible authorities approve the publication of this manuscript.

### Availability of data and material

Data and analysis scripts: https://doi.org/10.6084/m9.figshare.12798035.v1

### Code Availability

Code: https://github.com/AndrewBrughera/LSO_Transposed_Tones

### Author Contributions

Conceptualization: DM

Data Curation: AB

Formal Analysis: AB

Funding Acquisition: DM

Investigation: AB

Methodology: AB

Project administration: AB, JB, DM

Resources: DM, AB

Software: AB

Supervision: DM and JB

Visualization: AB

Validation: AB

Writing – Original Draft Preparation: AB

Writing – Review & Editing: AB, DM, JB

